# *Faecalibacterium prausnitzii* regulates carbohydrate metabolic functions of the gut microbiome in C57BL/6 mice

**DOI:** 10.1101/2024.09.30.615937

**Authors:** Peiling Geng, Ni Zhao, Yufan Zhou, Reuben S. Harris, Yong Ge

## Abstract

The probiotic impact of microbes on host metabolism and health depends on both host genetics and bacterial genomic variation. *Faecalibacterium prausnitzii* is the predominant human gut commensal emerging as a next-generation probiotic. Although this bacterium exhibits substantial intraspecies diversity, it is unclear whether genetically distinct *F. prausnitzii* strains might lead to functional differences in the gut microbiome. Here, we isolated and characterized a novel *F. prausnitzii* strain (UT1) that belongs to the most prevalent but underappreciated phylogenetic clade in the global human population. Genome analysis showed that this butyrate-producing isolate carries multiple putative mobile genetic elements, a clade-specific defense system, and a range of carbohydrate catabolic enzymes. Multiomic approaches were used to profile the impact of UT1 on the gut microbiome and associated metabolic activity of C57BL/6 mice at homeostasis. Both 16S rRNA and metagenomic sequencing demonstrated that oral administration of UT1 resulted in profound microbial compositional changes including a significant enrichment of *Lactobacillus*, *Bifidobacterium*, and *Turicibacter*. Functional profiling of the fecal metagenomes revealed a markedly higher abundance of carbohydrate-active enzymes (CAZymes) in UT1-gavaged mice. Accordingly, UT1-conditioned gut microbiota possessed the elevated capability of utilizing starch *in vitro* and exhibited a lower availability of microbiota-accessible carbohydrates in the feces. Further analysis uncovered a functional network wherein UT1 reduced the abundance of mucin-degrading CAZymes and microbes, which correlated with a concomitant reduction of mucin glycans in the gut. Collectively, our results reveal a crucial role of UT1 in facilitating the carbohydrate metabolism of the gut microbiome and expand our understanding of the genetic and phenotypic diversity of *F. prausnitzii*.

## Introduction

The human gastrointestinal tract is host to a diverse and complex community of microorganisms that influence host health and disease ^1–3^. These microbes, mainly anaerobic bacteria, encode a large variety of carbohydrate-active enzymes (CAZymes), including glycoside hydrolases, polysaccharide lyases, and carbohydrate esterases, and operate on a diverse range of dietary and host-derived substrates ^4^. The breakdown of complex carbohydrates not only fuels the bacterial needs but also produces bioactive compounds, particularly short-chain fatty acids (SCFAs), which have a profound impact on host epigenetic remodeling, energy expenditure, immune, and metabolic functions ^5^.

One of the most critical members of human gut microbiota is *Faecalibacterium prausnitzii*, representing 5 – 15% of the total gut bacterial population ^6^. *F. prausnitzii* is a major butyrate producer in the human gut and potentially the most important one ^6^. Oral administration of live *F. prausnitzii* or its supernatants is capable of reducing inflammatory responses in mice ^7–9^, and a reduction in *F. prausnitzii* abundance has been associated with a higher risk of several human intestinal ^10–12^, neurological ^13, 14^, and cardiovascular diseases ^15^ as well as type 2 diabetes ^16, 17^. Emerging evidence suggests that the functional properties of this bacterium are unlikely to be restricted to its production of butyrate ^9, 18, 19^. For instance, a 15-KDa protein, found in the supernatants of *F. prausnitzii* cultures, also demonstrates anti-inflammatory activity and thus protects mice against chemically induced colitis ^19^. In addition, *F. prausnitzii* produces formate and D-lactate, as two major fermentation end products^20^, and therefore may metabolically support members of gut microbiome community through bacterial cross-feeding. However, it is unclear whether different *F. prausnitzii* strains have specific functional determinants and employ differential mechanisms for intestinal function.

Earlier 16S rRNA-based comparison proposes the classification of *F. prausnitzii* strains into 2 phylogroups (I and II), whereupon ATCC 27768 and A2-165 are designated as the type strains of phylogroup-I and phylogroup-II, respectively ^21^. Subsequent genomic analysis leads to the identification of several distinct intraspecies phylotypes that harbor unique defense systems and carbohydrate-metabolizing capacity ^22, 23^. Recent meta-analyses of human gut metagenomes reveal at least five *F. prausnitzii* phylogenetic clades ^24, 25^, in which clade A is the most prevalent clade across global populations, detected in 78% of the human subjects ^25^. Clade-specific divergences may be driven by environmental clues and reflect their functional specifications ^24^. The prevalence and abundance of *F. prausnitzii* varies with age, geographical origin, and lifestyle. As such, newborns and elders have lower abundance compared to adults, whilst non-Western populations have significantly higher abundance than Western populations ^25^. Most recent studies suggest that *F. prausnitzii* strains also exhibit differential abundances in antibiotic resistance genes, virulence factors, and pathogenic genes ^26^ and that a single housekeeping gene may be used for qPCR-based quantification and classification ^27^.

Genomic diversity at sub-species level may lead to differences in phenotypes and metabolic functions. Indeed, specific gene orthologs in certain *F. prausnitzii* strains have been found at higher abundance in healthy individuals when compared to patients with mild cognitive impairment ^14^. Intraspecies compositional changes in *F. prausnitzii* may also occur in patients with Alzheimer’s disease, differentially correlated with the disease outcomes ^28^. Furthermore, strain-specific anti-inflammatory functions have been reported for the core *Faecalibacterium* amplicon sequence variant within the healthy Korean cohort ^29^. Thus, isolating and studying *F. prausnitzii* strains that represent different clades should be critical to understand their diversified functions in the gut. However, most of the animal studies have been conducted with *F. prausnitzii* strain A2-165 (clade C) ^9, 18, 30–33^, and much less is known so far regarding representatives of other clades, particularly the most prevalent clade A. Furthermore, numerous studies have examined the correlation between *F. prausnitzii* and intestinal disorders, and the beneficial properties of this bacterium have been largely attributed to its production of butyrate ^6, 34, 35^. However, fewer studies have examined the direct impact of *F. prausnitzii* on the gut microbiome ^7, 8, 30, 33^ and particularly associated metabolome at homeostasis. Here, we show that *F. prausnitzii* strain UT1 (abbreviated as UT1 hereafter), a new *bona fide* member of the clade A, modifies the gut microbiota and associated carbohydrate profiles and controls the abundance and activity of mucin-degrading microbes. Our findings provide insights into the role of *F. prausnitzii* in the regulation of gut microbiome and may contribute to designing probiotic strategies for the improvement of intestinal barrier function.

## Results

### Isolation and identification of a novel *F. prausnitzii* strain that belongs to a unique phylogenetic clade

*F. prausnitzii* exhibits substantial intraspecies diversity that is associated with disease conditions^25^. To elucidate the functional diversity of *F. prausnitzii*, efforts were made to isolate bacterial strains phylogenetically distinct from the type strains (A2-165 and ATCC 27768). Using strict anaerobic culturing and rapid PCR screening, a new strain, named <u>U</u>niversity of <u>T</u>exas 1 (UT1), was isolated from the stools of heathy human donor. *De novo* assembly using Oxford nanopore long reads produced the complete circular genome of UT1 bacterium, which was further polished (error correction) using high-quality Illumina short reads. Both sequencing approaches resulted in average coverages of >2000x. Comparative genome analysis demonstrated significant sequence diversity to available *F. prausnitzii* genomes, including a predicted prophage uniquely presented on the genome of UT1 rather than other strains (**Figure 1A**). This prophage carries genes related to phage DNA replication and packaging, morphogenesis, and host cell lysis (**Figure S1A**). A BLAST search revealed >96% sequence identity to bacteriophages derived from human fecal metagenomes, suggesting its prevalence and potential role in driving horizontal gene transfer in human gut.

**Figure 1.**
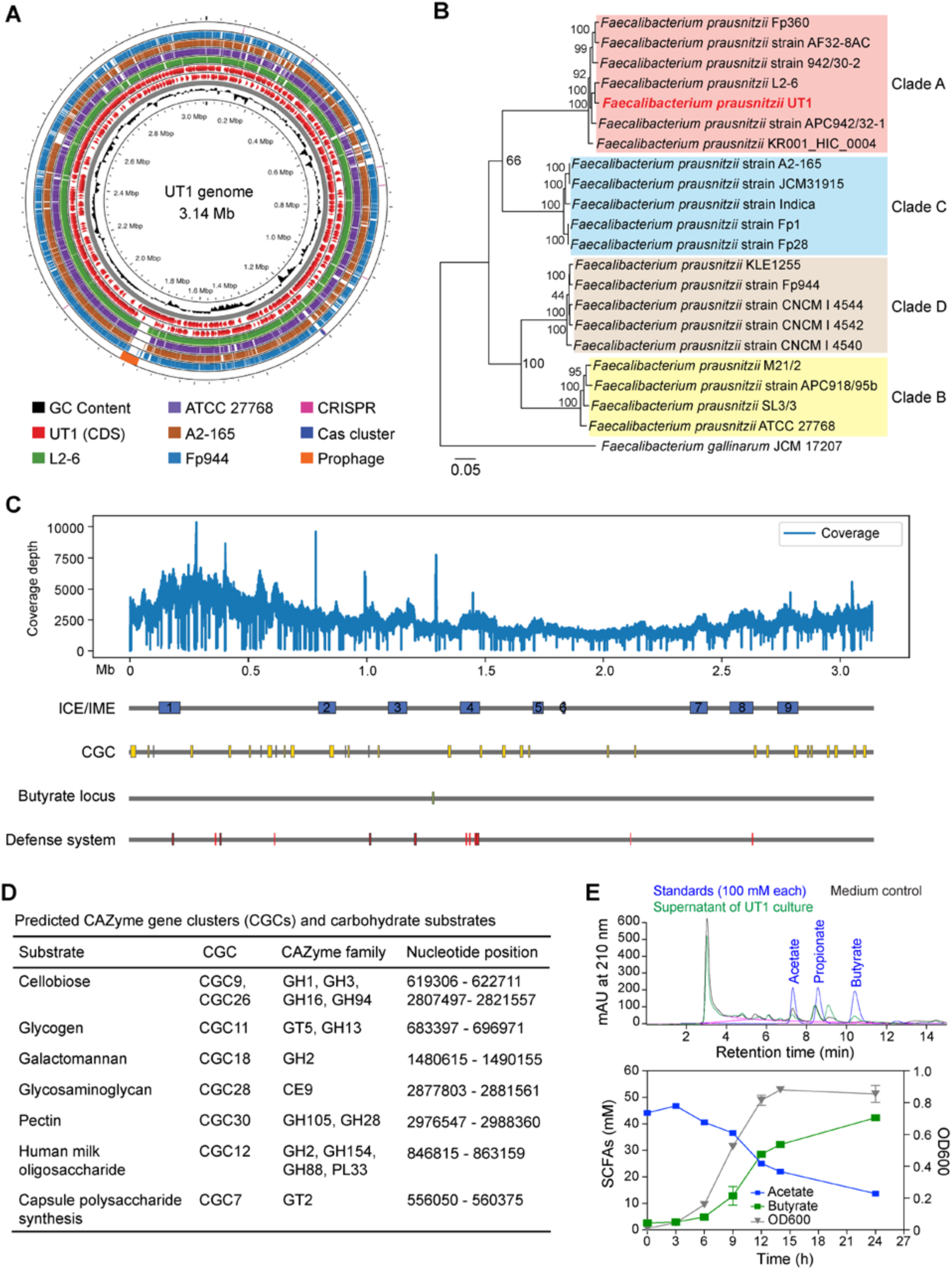
Genomic analysis of UT1. **(A)** Genome comparison of UT1 with representative strains of *F. prausnitzii* clades A-D. Rings with different colors indicate different bacterial genomes. CRISPR-Cas system and putative prophage were shown in the outermost ring. UT1 was used as reference, and CDSs with opposite orientations were depicted in two separate rings (red). **(B)** Whole genome phylogenetic analysis of 22 *Faecalibacterium* isolates. The tree was built on the protein and gene sequences for 100 single-copy genes found in all the selected genomes. Brach support values are obtained using 100 bootstrap and the length of a branch is relative to the total number of changes at each site. **(C)** Coverage plot of UT1 genome sequencing reads and locations of integrative and conjugative/mobilizable elements (ICE/IMEs), CAZyme gene clusters (CGCs), butyrate locus and defense systems. **(D)** dbCAN3 analysis of CAZyme gene clusters (CGCs) and their carbohydrate substrates. **(E)** HPLC chromatograms of distinct acetate and butyrate peaks in the UT1 culture compared to standards and the medium controls (top panel). Levels of acetate and butyrate in the supernatants of UT1 culture (n = 3 samples/time point) are shown in the bottom panel.

Whole genome phylogenetic analysis showed UT1 was a *bona fide* member of the clade A, whereas ATCC 27768 and A2-165 were each represented by a different clade (B and C, respectively) (**Figure 1B**). Further analysis identified 9 integrative and conjugative/mobilizable elements (ICE/IMEs) that were scattered across the whole genome (**Figure 1C**). Of these, ICE/IME-2 expresses the components of human milk oligosaccharide metabolism and type IV secretion systems important for interbacterial DNA transfer (**Figure 1C and Figure S1A**). Several defense systems, including restriction-modification systems, were also detected within the ICE/IMEs (**Figure 1C**). Additionally, a conserved type I-C CRISPR-Cas system was found in strains of clade A, but not other clades, which instead possess a type I-E system (**Figure S1B**), implying clade-specific defense mechanisms.

*F. prausnitzii* possesses the capability to metabolize a wide range of complex carbohydrates and different clades may have specific carbohydrate utilization patterns ^21, 25, 36, 37^. To gain insight into the carbohydrate-metabolic potential of UT1, we analyzed the CAZymes and CAZyme gene clusters (CGCs), which are experimentally characterized polysaccharide utilization loci ^38^, and predicted their substrates. There was a total of 32 CGCs distributed across the genome (**Figure 1C**), among which 7 CGCs were predicted to metabolize specific substrates, including cellobiose and pectin. Interestingly, 2 of the CGCs, potentially catabolizing human milk oligosaccharide (CGC12) and cellobiose (CGC26) (**Figure 1D**), were found on the ICE/IMEs (**Figure 1C**). By comparison of CGCs obtained from strains of different clades, we found the CGC responsible for galactomannan utilization was intact in clade A, whereas the involved carbohydrate transporters were lacking in strains of clades B, C or D (**Figure S1C**).

The major end products of the microbial carbohydrate-fermenting activity in the gut are SCFAs – in particular butyrate ^5^. To examine the butyrate-producing capacity of UT1, we searched its whole genome and identified a butyrate locus (**Figure 1C**). High-performance liquid chromatography (HPLC) analysis demonstrated that UT1 abundantly synthesized butyrate in growth medium in the presence of soluble starch, and the production was gradually increased along with bacterial growth, whereas the levels of acetate were proportionally deceased (**Figure 1E**). These results suggest that UT1 is equipped with an active molecular machinery for butyrate biosynthesis.

### Regulation of gut microbiome composition

Commensal microbes play a critical role in shaping gut microbiota and metabolic functions that are specific to bacterial strains ^39^. To elucidate the intestinal microbial responses to UT1, we employed multiomic approaches to comprehensively profile the impacts of UT1 on gut microbiome and associated metabolome (**Figure 2A**). To proceed, conventionally housed C57BL/6 mice were gavaged with UT1 or PBS every 2 days for a total of 7 gavages, and fecal samples were collected on day 14 to analyze the gut microbiome by 16S rRNA sequencing. While there was no difference in alpha-diversity (**Figure S2A**), administration of UT1 resulted in a significant shift in the overall microbial community (PERMANOVA = 0.006), as demonstrated by a weighted UniFrac principal coordinate analysis (PCoA) (**Figure 2B**). The relative abundance of the phylum Actinobacteria was increased about 3.5-fold in UT1-gavaged mice compared to PBS controls (**Figure 2C**), which was due to elevated class Actinobacteria and marginally increased Coriobacteriia (**Figure 2D**). Major compositional alterations were seen within the phylum Firmicutes, where the abundance of the class Clostridia was significantly reduced from 31.6% to 19.1%, and the levels of Bacilli were increased from 11.6% in PBS group to 23.7% in UT1 group (**Figure 2D**). Accordingly, Lactobacillaceae (class Bacilli) and Turicibacteraceae (class Erysipelotrichia), as two major families of Bacilli, were notably enriched in UT1 group compared to PBS controls (**Figure 2E**). Bacterial family changes were also observed within Clostridia. Specifically, Lachnospiraceae, Ruminococcaceae and undesignated/unclassified Clostridiales were all reduced in UT1-treated mice. In contrast, Clostridiaceae, which was detected at marginal level in control mice, expanded massively after 2 weeks of UT1 gavage, accounting for 4.1% of total bacteria. Additionally, the relative abundance of Bifidobacteriaceae, a generally considered probiotic and health-promoting bacterial family, was also elevated in UT1-treated mice (**Figure 2E**).

**Figure 2.**
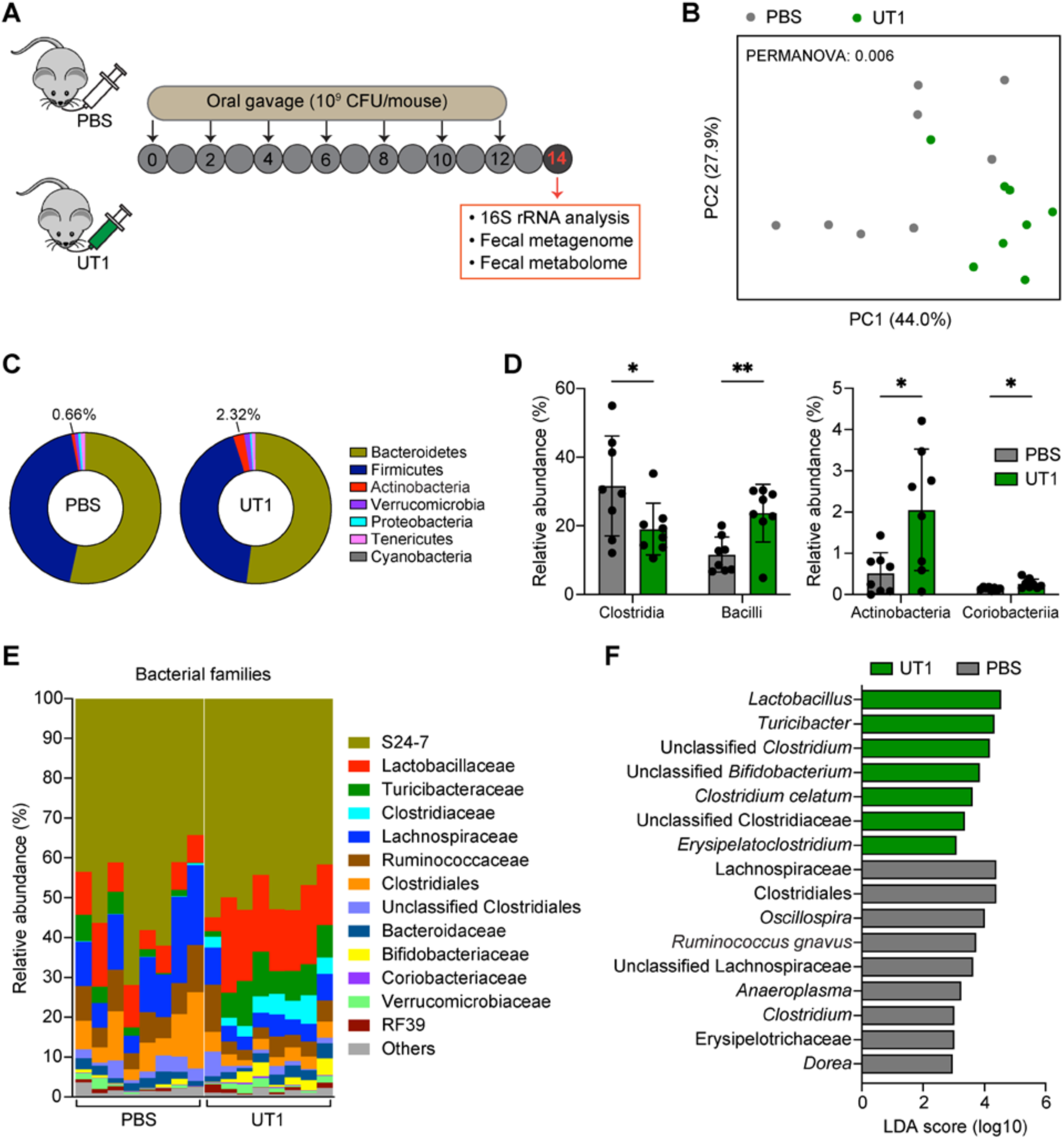
Gut microbiome analysis of mice treated with UT1. **(A)** Experimental scheme. Groups of C57BL/6 mice (7–8 weeks old) were orally gavaged with UT1 (10^9^ CFU/mouse) or PBS every 2 days. Feces were collected from the two groups of mice on day 14 for 16S rRNA, metagenomics, and metabolomics analyses. **(B)** Weighted UniFrac principal coordinate analysis plot depicts the bacterial community structures (n = 8/group). **(C)** Pie charts showing the bacterial phylum distribution. **(D)** Top bacterial class abundance. **(E)** The relative abundance of top bacterial families. **(F)** LEfSe plot showing differentiating taxa (species level) between UT1- and PBS- gavaged mice. *p < 0.05 and **p < 0.01, unpaired student *t*-test.

Linear discriminant analysis effect size (LEfSe) was conducted to explore bacterial taxa at species level that explain the majority of the differences between the two groups. Here, *Lactobacillus*, *Turicibacter* and *Bifidobacterium*, critically involved in regulation of host metabolism contributing to intestinal health ^40, 41^, were the top taxa enriched in UT1 group (**Figure 2F**). *Clostridium celatum*, which possesses glycosyl hydrolases specific to mucins ^42^, was also increased in these mice. In contrast, UT1 majorly restricted the colonization of Clostridiales members, including Lachnospiraceae, reported to be associated with worsened recovery after surgery for Crohn’s disease ^43^. *Ruminococcus gnavus*, populations of which bloom during flares of symptoms in patients with Crohn’s disease and lupus nephritis ^44, 45^, and *Dorea*, which has been identified as a microbiome signature of pediatric liver disease ^46^, were also reduced in mice receiving UT1. Both *Ruminococcus gnavus* and *Dorea* have been also reported to degrade mucins and utilize mucin-derived glycans for their growth ^47, 48^, illustrating the potential influence on mucin-degrading capacity of the microbiome.

*Oscillospira* is positively associated with leanness and health ^49^, however, higher numbers of *Oscillospira* are also found in patients with impaired intestinal function such as constipation ^50^. Diminished levels of *Oscillospira* were found in UT1-gavaged mice, which also exhibited the decreased abundance of Erysipelotrichaceae (**Figure 2F**), found to be enriched in IBD patients and animals, albeit with conflicting findings ^51^. These results indicate an obvious role of UT1 regulating gut microbiome structure and composition.

### Microbial signatures revealed by metagenomics analysis

To further determine the compositional changes of gut microbiota at a high-resolution level, we performed metagenomics sequencing of fecal samples collected from UT1- and PBS-gavaged mice (**Figure 2A**). After removal of sequence reads mapping to the mouse genome, the microbiome sequence reads (>50 million paired-end reads/sample) were mapped to ∼1 million microbial genomes to predict the abundance of microorganisms. Consistent to 16S rRNA analysis, no difference in Shannon index was observed. However, there was a slight increase in bacterial richness in UT1 group (**Figure 3A**). An estimation of the beta-diversity demonstrated a clear separation of microbial communities between the two groups of mice (**Figure 3B**). Metagenomic sequencing identified differential bacterial families that were largely consistent with 16S rRNA sequencing, including reduced levels of Lachnospiraceae and increased abundance of Lactobacillaceae, Bifidobacteriaceae and Turicibacteraceae in the UT1 group compared to the PBS control (**Figure 3C**). It is noted that a lower abundance of Lactobacillaceae and a higher abundance of Clostridiaceae were detected by metagenomics analysis compared to 16S rRNA analysis in the two cohorts of PBS mice (**Figure S2B**), which could be due to variations in sample preparation and data analysis ^52^.

**Figure 3.**
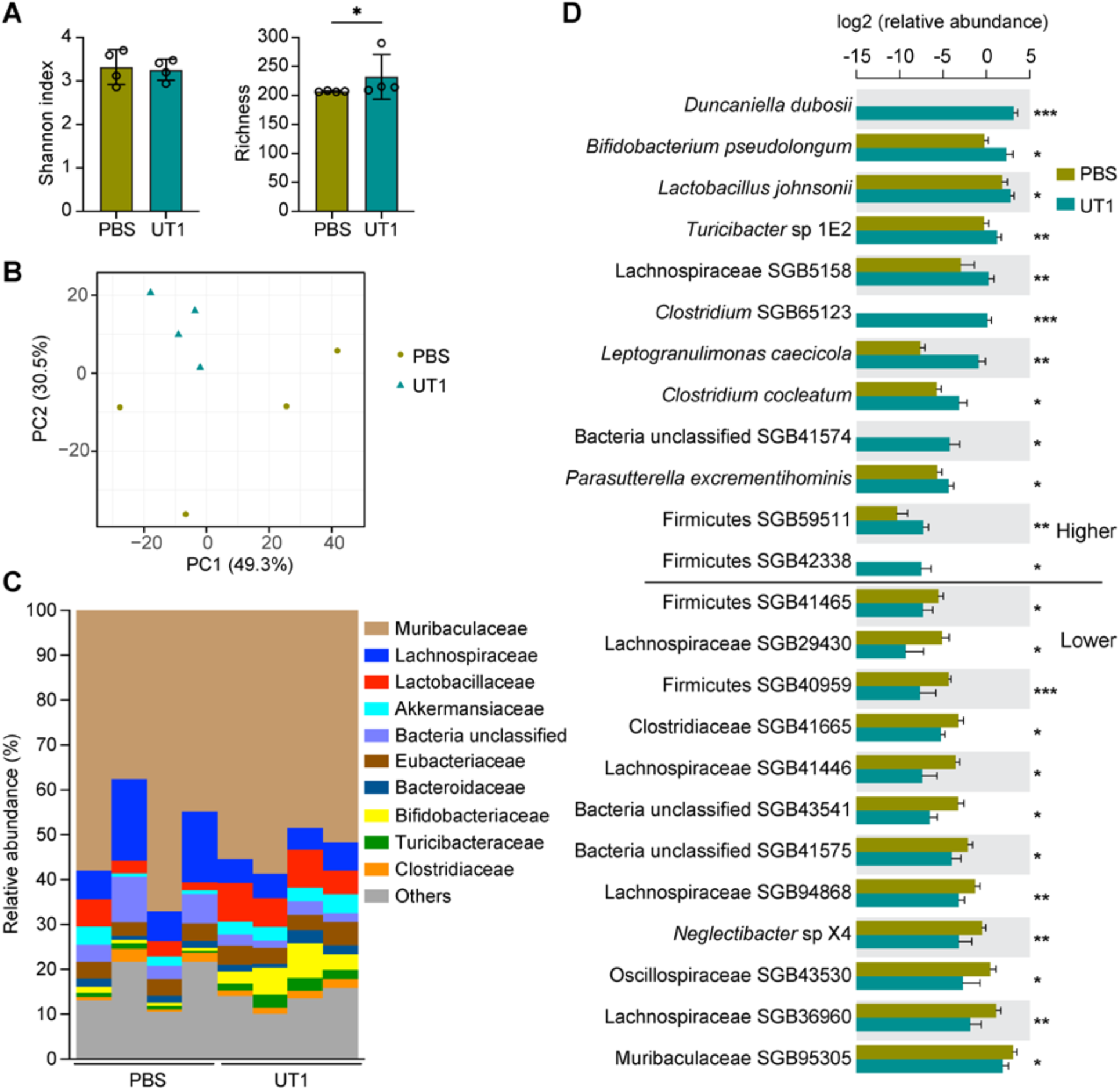
Microbial signatures revealed by metagenomics sequencing. **(A)** Alpha-diversity of the fecal metagenomes in UT1- and PBS-gavaged mice (n = 4/group). **(B)** Bacterial community structures of the two groups of mice. **(C)** Top bacterial family abundance. **(D)** Differentially enriched bacterial species between UT1- and PBS-gavaged mice. SGB, species-level genome bin. *p < 0.05 and **p < 0.01, unpaired student *t*-test.

Our further compositional analysis was confined to the 258 unique metagenomic species, resulting in the identification of 24 differentially enriched species-level genome bins (SGBs), most of which are yet-to-be-characterized species (**Figure 3D**). *Duncaniella dubosii* (family Muribaculaceae/S24-7), which has recently been shown to protect mice against chemical-induced colitis ^53^, was highly enriched in the feces of UT1-gavaged mice (9.0% of total bacteria), in striking contrast to its undetectable levels in control group (**Figure 3D**). The increased relative abundances of *Leptogranulimonas caecicola* and *Parasutterella excrementihominis* (**Figure 3D**), which are implicated in bile acid metabolism ^54, 55^, were also observed. In accordance with 16S rRNA profiling (**Figure 2F**), metagenomics analysis further demonstrated the significant enrichment of specific bacterial species, including *Lactobacillus johnsonii*, *Bifidobacterium pseudolongum* and *Turicibacter* sp 1E2 (**Figure 3D**). Similarly, the reduced species in UT1 group were primarily derived from Clostridia, such as SGBs of Lachnospiraceae, Oscillospiraceae, and Clostridiaceae, once again illuminating a UT1-dependent microbial signature.

### CAZyme abundance and diversity of the gut microbiome

Next, we analyzed the abundance of CAZymes and predicted their substrates in the fecal metagenomes of UT1- and PBS-treated mice using dbCAN3 ^38^. We detected 47 differentially enriched CAZymes (15.0% of total CAZymes), 37 of which had higher abundance in mice gavaged with UT1 compared to PBS (**Figure 4A**). Specifically, 8 glycoside hydrolases (e.g., glucosidase, fucosidase, and rhamnosidase) and 2 phosphorylases (maltose and sucrose phosphorylases), which catalyze the liberation of hexoses from carbohydrate polymers, were enriched in UT1 group (**Figure 4B**). Pectinesterase and xanthan lyase, involved in the breakdown of pectin and xanthan, were also enriched. CAZymes for synthesizing bacterial cell wall components, including peptidoglycan glycosyltransferase and diacylglycerol 3- glucosyltransferase, potentially related to the turnover of microbes, were also found at higher levels in UT1-gavaged mice. Intriguingly, we detected markedly altered abundance of CAZymes related to mucin degradation (**Figure 4C**), including the reductions in GH16_21 that may cleave the poly-N-acetyllactosamine ^4^, GH89 and GH129 that remove galactose and its derivative N-acetyl-galactosamine, and a slight increase in GH29 for the removal of fucose residues ^56^, suggesting a modification in microbial genetic potentials to degrade mucins.

**Figure 4.**
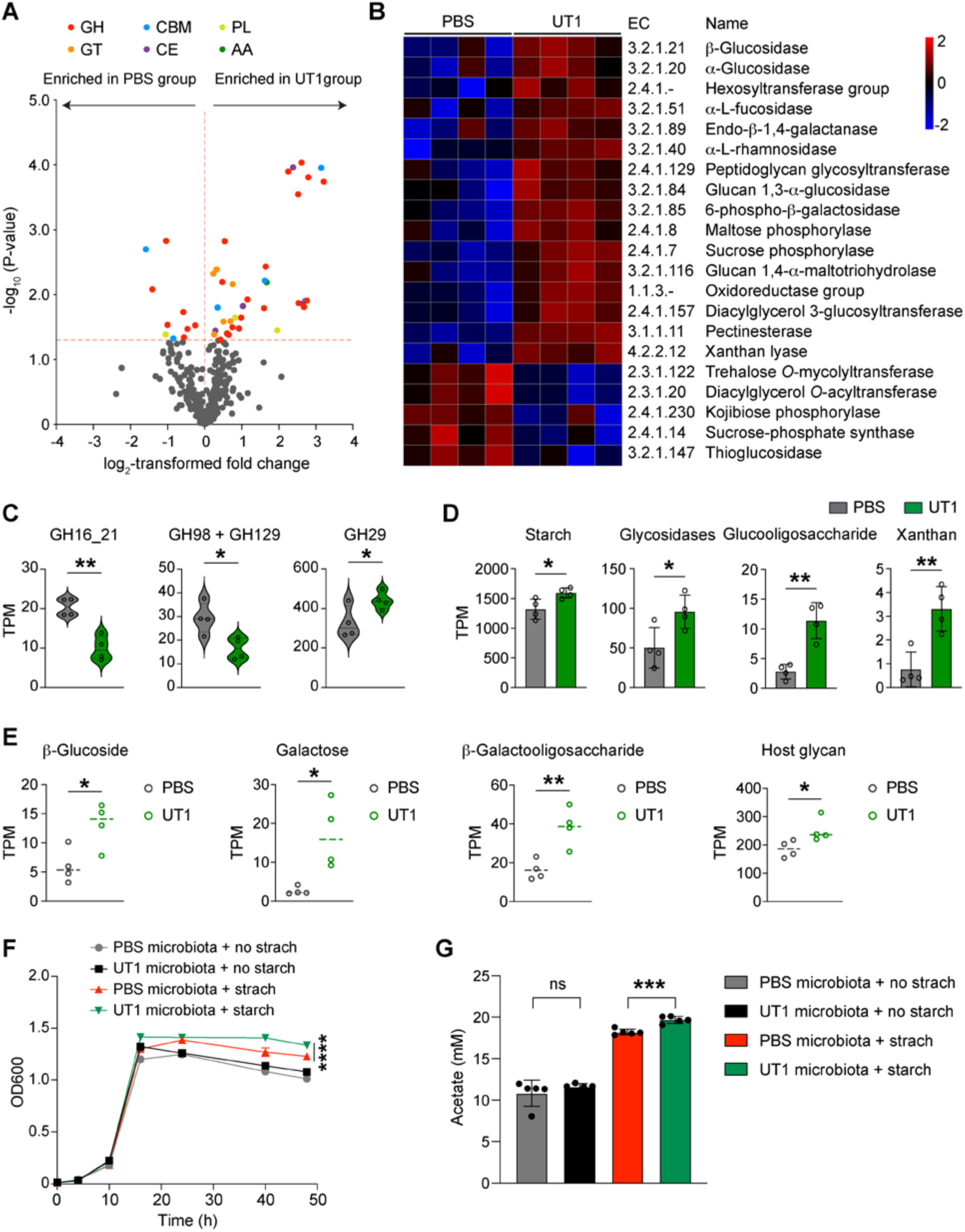
CAZyme abundance and diversity of the fecal metagenomes. **(A)** Volcano plot of CAZymes differentially enriched in UT1- and PBS-gavaged mice (n=4/group). GH, glycoside hydrolase; GT, glycosyltransferase; CBM, carbohydrate-binding module; CE, carbohydrate esterase; PL, polysaccharide lyase; AA, enzymes of auxiliary activities. **(B)** Heatmap of CAZyme ECs with significantly differing abundance. **(C)** The abundance of CAZymes related to mucin-degrading. TPM (Transcripts Per Million) is used as a measurement of the normalized CAZyme abundance. **(D)** The total abundance of CAZymes predicted to utilize indicated substrates. **(E)** The abundance of carbohydrate-utilizing CGCs in the two groups of metagenomes. **(F)** Feces were collected from UT1- (n=9) and PBS-gavaged mice (n=5), and pooled to generate fecal microbiota suspension. The *in vitro* anaerobic growth of the UT1-conditioned microbiota versus PBS control microbiota in the presence or absence of starch (n=5 cultures/group). **(G)** HPLC measurement of acetate production by UT1 microbiota and PBS microbiota after growth in the gut microbiota culture medium (GMM) for 48 hours. Data are presented as mean ± SD. *p < 0.05, **p < 0.01, ***p < 0.001, ****p < 0.0001, unpaired student *t*-test.

The dbCAN-sub search and substrate mapping revealed the increased abundance of CAZymes responsible for fermentation of complex carbohydrates such as starch and xanthan (**Figure 4D**). Furthermore, we performed dbCAN-PUL search against the metagenomes at CGC levels to estimate the abundances of polysaccharide utilization loci and predict glycan substrates. The resulting CGCs, implicated in the metabolism of glucoside, galactose, β-galactooligosaccharide, and host glycan, were all enriched in UT1-gavaged group (**Figure 4E**).

To investigate if enriched CAZymes abundances in fecal metagenomes could lead to elevated capability of the microbiota to metabolize carbohydrates such as starch, we cultured the fecal microbiotas of UT1- and PBS-gavaged mice in a gut microbiota culture medium (GMM) supplemented with or without starch. The bacterial growth and acetate production, as a measurement of metabolic activity, were examined in the anaerobic culture media. Here, no obvious differences were detected between the two groups of cultured fecal microbiotas when grown in GMM without starch (**Figure 4, F and G**). However, the bacterial growth and acetate production were notably stimulated upon adding starch in the GMM, and the cultured UT1 microbiota had better growth and higher acetate release compared to the control microbiota (**Figure 4, F and G)**, indicating the enhanced carbohydrate utilization.

### Programming of microbiota-associated metabolome

To further elaborate on the functional changes of the gut microbiome, we conducted untargeted metabolomics analysis on the fecal samples collected from the same UT1- and PBS-treated animals in metagenomics experiments (**Figure 2A**). High-resolution liquid chromatography-mass spectrometry (LC-MS) analysis demonstrated that oral administration of UT1 led to a noticeable shift in the global metabolomic profiles of the fecal microbiota, as revealed by principal component analysis (PCA) of metabolite features identified by both positive and negative ionizations (**Figure 5A**). Carbohydrate metabolic pathways, including N-glycan degradation and galactose metabolism, were the top pathways differentially enriched between the fecal samples of UT1- and PBS-gavaged mice (**Figure 5B**). Glycolysis and gluconeogenesis, pyruvate metabolism and pentose phosphate pathways, which are all involved in the catabolism of glucose, as well as pathways related to amino acids (e.g., histidine, arginine and proline), were also significantly modified by UT1. By comparing to respective standards, 32 metabolites were identified with intensities significantly differing between the two groups (**Figure 5C**). Here, lactic acid, abundantly produced by lactic acid bacteria such *Lactobacillus* and *Bifidobacterium*, was increased in UT1-gavaged mice, which also had increased N-acetylneuraminate and D-alanyl-D-alanine, two major constituents of the bacterial peptidoglycans. N-acetylneuraminate is also the predominant sialic acid in human and mammals that exhibit antioxidative activities ^57^. Consistently, a panel of antioxidant metabolites, particularly putrescine and alanine, was detected with elevated amounts in mice gavaged with UT1. Additionally, N-acetyl-L-aspartate, which may inhibit glycation inducing protein denaturing and oxidative stress ^58^, was also enriched in these mice (**Figure 5C**). Oxidation of polyunsaturated fatty acids such as linoleic acid generates oxidative stress and thus has been linked to various pathological diseases ^59^. While the level of linoleic acid was maintained in UT1-gavaged mice, the medium-chain saturated fatty acids (azelaic acid and suberic acid), which can be generated from the oxidation of linoleic acid, were significantly reduced, further indicating the active control of oxidative reactions in the gut.

**Figure 5.**
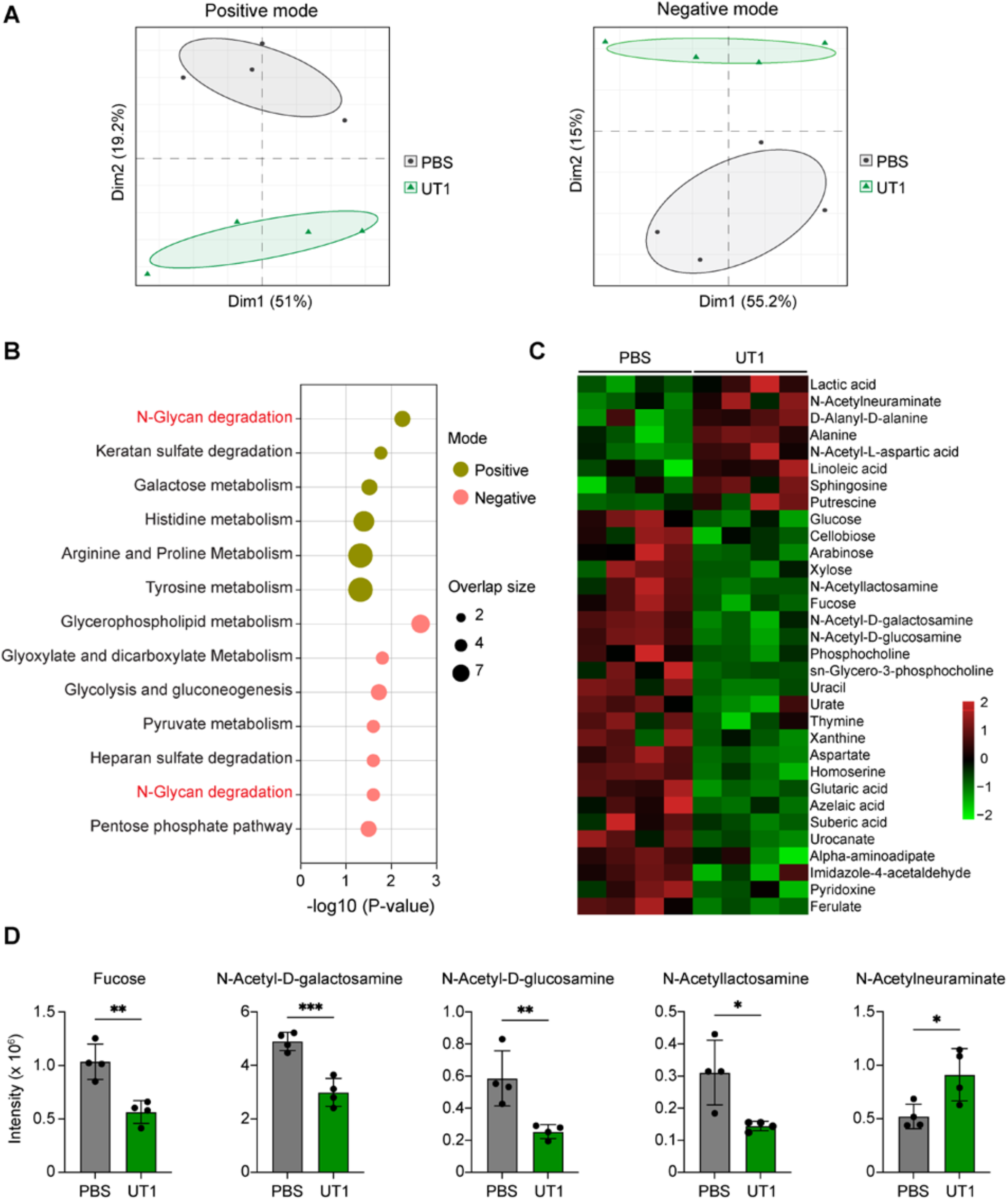
Metabolomic profiles of fecal microbiota. **(A)** Principal component analysis (PCA) plots of metabolite features identified by positive and negative ionizations in the feces of UT1- and PBS-gavaged mice (n = 4/group). **(B)** Metabolic pathway analysis of metabolites with significantly differing intensity between UT1 and PBS groups. The overlapped size indicates the number of significant metabolic features mapped to corresponding pathways. **(C)** Heatmap showing differentially enriched metabolites. **(D)** Bar graphs depicting the intensities of mucin-related glycans. Note: LC-MS analysis was repeated twice on the same fecal samples, and similar results were obtained. *p < 0.05, **p < 0.01, ***p < 0.001, unpaired student *t*-test.

Moreover, the metabolites with reduced intensities in UT1 group were primarily defined by a group of microbiota-accessible carbohydrates (e.g., arabinose, xylose, and cellobiose) (**Figure 5C**), along with enriched CAZymes for carbohydrate breakdown (**Figure 4**), suggesting a critical role of UT1 in supporting microbial carbohydrate metabolism. Interestingly, the mucin-decorating glycans, particularly N-acetyl-D-galactosamine, N-acetyl-D-glucosamine and N- acetyllactosamine, which are cleaved from mucins through specific microbial CAZymes (**Figure 4C**), were also highly decreased (**Figure 5D**), indicative of controlled mucin-degrading activities in UT1-treated mice. In addition, several nucleobases (e.g., uracil and thymidine) and amino acids (e.g., aspartate) as well as metabolites associated with lysine degradation (alpha-aminoadipate) and histidine metabolism (urocanate and imidazole-4-acetaldehyde), were detected at lower levels in UT1 group than the control group (**Figure 5C**), documenting an unneglectable impact on microbial metabolic function.

### Correlation analysis of gut microbiota and associated metabolites

Next, we explored the functional link between the gut microbiome and fecal metabolomes of the same animals by calculating the Spearman’s correlation coefficients. Significant microbe-metabolite correlations (|*r*| > 0.85, FDR < 0.05) were identified between 24 differentially enriched metagenomic species and 32 significantly altered metabolites (**Figure 6**). Apparently, *Duncaniella dubosii* showed negative correlation with the carbohydrates, mainly glucose, arabinose and fucose, whereas *Parasutterella excrementihominis*, *Turicibacter* sp 1E2 and *Bifidobacterium pseudolongum* demonstrated positive association with antioxidant alanine and N-acetyl-L-aspartic acid. Lachnospiraceae SGB41446, *Neglectibacter* sp X4 and Clostridiaceae SGB41665, all of which had reduced abundance in UT1 group, were negatively associated with linoleic acid and positively associated with suberic acid or azelaic acid, illustrating a possible role of these bacteria in fatty acid oxidation. While the mucin-related fucose and N-acetyllactosamine were positively associated with a species of Lachnospiraceae (SGB36960), N-acetyl-D-galactosamine and N-acetyl-D-glucosamine were reversely correlated with *Leptogranulimonas caecicola.* Additionally, there was one species within Gram-positive Firmicutes (SGB59511) that showed significant correlation with N-acetylneuraminate (**Figure 6**). Together, these results correlatively illuminate a crucial role of UT1 in regulating gut microbiome and associated metabolic activities that may control the oxidative stress and barrier dysfunction in the gut.

**Figure 6.**
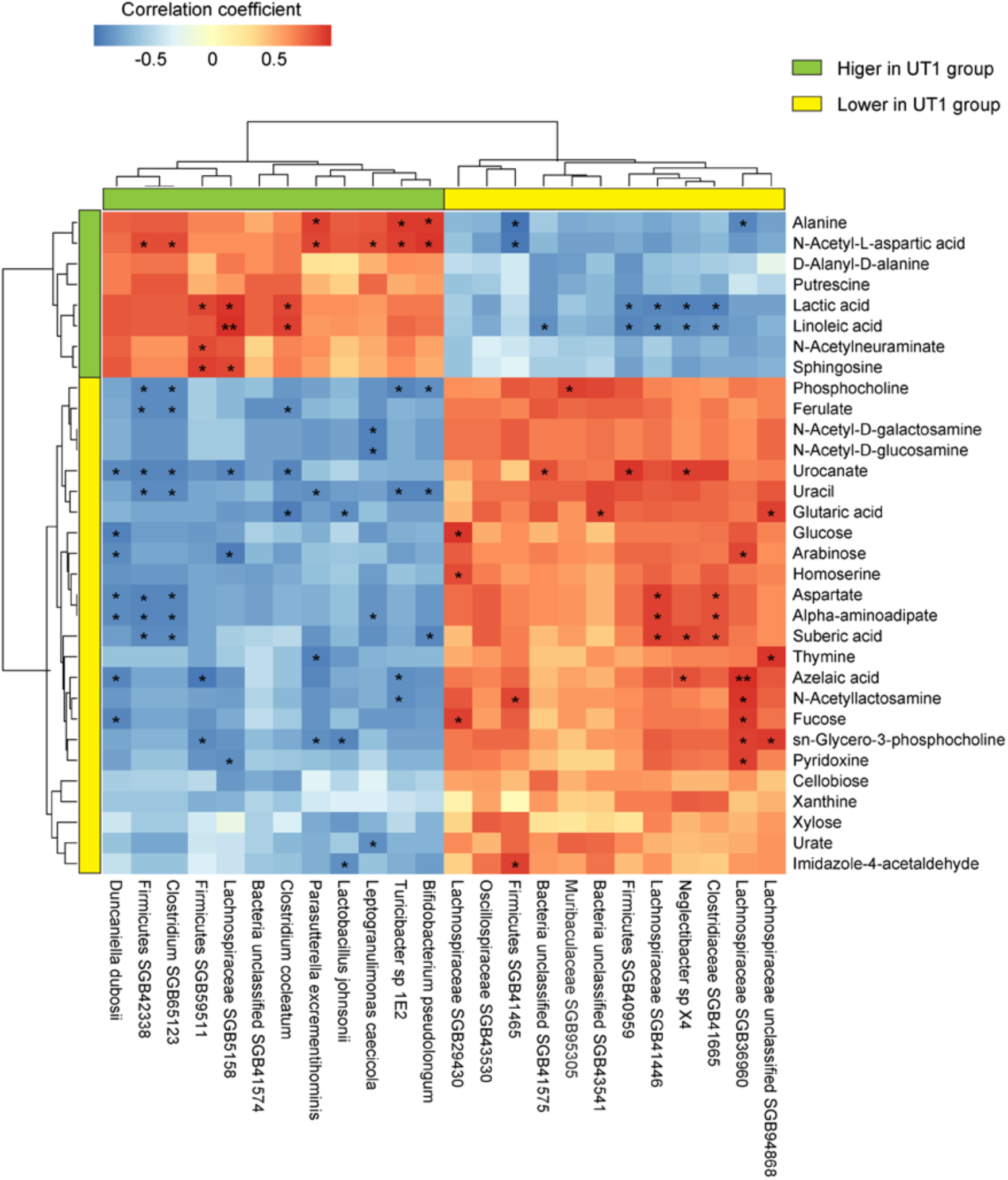
Correlation analysis of microbiome and associated metabolites. The correlations between significant metagenomic species and differentially enriched metabolites were determined by calculating Spearman’s correlation coefficients. Positive and negative correlations are shown as red and blue in the heat map, respectively. Significant microbiota-metabolite correlations were determined based on the criteria (*r* > 0.85 and FDR < 0.05). *FDR < 0.05 and **FDR < 0.01.

## Discussion

Commensal bacteria contribute to intestinal hemostasis mainly through the production of essential metabolites or binding to their cognate receptors that translate the nature of the microorganisms into immunogenic or immunotolerant cell programs ^60, 61^. Impairment of microbial metabolic activity or disruption of the balanced microbe-host communications, as a result of dysbiosis, may result in pathogenic inflammation inducing intestinal tissue damage. Thus, understanding how gut commensal bacteria, particularly those dominantly occupying the gastrointestinal tract, contributes to the modulation of intestinal microbiome and metabolomic homeostasis is of paramount importance. In this study, we characterized a novel gut commensal strain, representing the most prevalent phylogenetic clade of *F. prausnitzii* family, and demonstrated its essential role in the regulation of microbiome carbohydrate metabolism. To the best of our knowledge, this is the first study to comprehensively examine the impacts of *F. prausnitzii* on the fecal microbiome by integrating 16S rRNA, metagenomics, and metabolomics analyses at homeostatic condition.

Several *bifidobacterium* strains, including *B. calenulatum* and *B. adolescentis*, have been shown to promote the production of butyrate by *F. prausnitzii* ^62, 63^. Human microbiota studies also suggest the concurrent increase of *B. adolescentis* and *F. prausnitzii* upon consuming complex carbohydrates ^64^. However, little evidence supports the bifidogenic effects of *F. prausnitzii*. We clearly showed the significant enrichment of *B. pseudolongum* in the feces after UT1 treatment, further emphasizing the mutualistic cross-feeding between *F. prausnitzii* and *Bifidobacterium in vivo*. Additionally, we detected higher levels of *L. johnsonii*, which could confer numerous health benefits to the host ^65^, and correspondingly enriched lactic acid in the feces of UT1-gavaged mice. It is also noted that specific strains of *Lactobacillus* have been shown to elevate the abundance of *F. prausnitzii* in chickens ^66, 67^. In contrast, a recent human study document that the intake of *L. johnsonii* by healthy volunteers decreases the fecal levels of *F. prausnitzii* ^68^. This discrepancy could be due to the phenotypic diversity of specific bacterial strains, but could also vary with other features such as host genetics and baseline microbiota structures ^39^.

In rodent and human studies, *Turicibacter* has been shown to play a crucial role in modulating bile acid transformation and lipid profiles ^41, 69^, and its relative abundance often negatively correlates with host adiposity ^70^. Reduced adipose tissue inflammation and body weight gain have been also reported in *F. prausnitzii*-treated mice fed a high-fat-diet ^8, 71^. We observed a significant increase in *Turicibacter* and differential alterations in fatty acids in mice gavaged with UT1, leading to a possibility that *F. prausnitzii* may support the growth of *Turicibacter* to regulate host lipid metabolism. In addition, butyrate drives mitochondrial fatty acid β-oxidation in intestinal epithelial cells ^72^ and dietary supplementation of butyrate has been also shown to prevent high fat-diet induced obesity ^73^. Although UT1 highly produced butyrate in the culture, we failed to detect butyrate in the fecal samples, which could be due to reduced levels of microbes and in particular Clostridia that also synthesize this metabolite, as previously reported ^33^. However, it is also conceivable that genetically different *F. prausnitzii* strains may exhibit functional diversity ^23, 25^ resulting in differential regulation of the gut microbiome. Further studies are still required to elucidate the effects of UT1 and the associated metabolic regulation in the context of obesity.

A fundamental function of gut microbiota is to breakdown complex carbohydrates, providing bioactive metabolites and micronutrients that not only influence host metabolism but also reshape the gut microenvironment, synergistically contributing to the metabolic health of the host ^74^. The microbiota derived from UT1-treated mice apparently had elevated genetic capacity and heightened metabolic activities to degrade carbohydrates, as correlatively demonstrated by CAZyme mapping and metabolomics analysis. The enriched abundance of CAZymes and CGCs reflects the compositional changes of the microbiome and is unlikely attributable to the overrepresentation of UT1 DNA in the feces, as this bacterium was undetected by either 16S rRNA or metagenomics sequencing. Indeed, colonization experiment also showed that UT1 only transiently colonized the gut, and no UT1-specific DNA was detected 2 days after the gavage (**Figure S2C**). However, we cannot exclude the possibility that the bacterium may consume fecal carbohydrates during the transient colonization, thus contributing to the reduced bioavailability of carbohydrates in the feces.

In the gut, mucins are heavily glycosylated proteins produced by goblet cells and play a crucial role in intestinal barrier function, and reduced mucus layer has been associated with IBD and increased susceptibility to pathogens ^75^. The mucin glycans (e.g., N-acetyl-D-galactosamine) and associated CAZymes that catalyze the removal of these glycans from mucins were reduced in mice gavaged with UT1. Consistently, diminished levels of microbes that possess the genetic capability to degrade mucins, including *Dorea* and *Ruminococcus gnavus*, were also observed in these mice. *F. prausnitzii* also appears to attenuate *Bacteroides thetaiotaomicron*-induced mucin expression and glycosylation by consuming acetate, which may upregulate the expression of the transcription factor KLF4 essential for goblet cell differentiation ^76^. These results may suggest a general role of *F. prausnitzii* in regulating the glycosylation profiles of the mucins.

In summary, our results demonstrate a critical role of UT1 in coordinating intestinal microbiome composition and carbohydrate metabolic function, and further emphasize the importance of understanding the functional diversity of *F. prausnitzii*, which may be collectively considered to maximize their contribution to the health of the host.

## Materials and Methods

### Bacterial growth condition

Stool samples were obtained from a de-identified healthy human donor (female), diluted and plated on reinforced clostridial agar containing 10 g peptone, 10 g beef extract, 3 g yeast extract, 5 g dextrose, 1 g soluble starch, 5 g sodium chloride, 3 g sodium acetate, 0.5 cysteine hydrochloride, 0.5 mg resazurin, and 15 g agar per liter (pH 6.8). Single colonies were selected for PCR screening using *F. prausnitzii* 16S rRNA-specific primers (forward: 5’-GGAGGAAGAAGGTCTTCGG-3’; reverse: 5’-AATTCCGCCTACCTCTGCACT-3’; 248-bp). Bacteria were grown in reinforced clostridial medium (RCM, no agar) at 37°C in an anaerobe chamber (model AS-500, Anaerobe Systems) with a gas mixture of 90% nitrogen, 5% carbon dioxide, 5% hydrogen.

### Whole-genome sequencing

Complementary Oxford Nanopore and Illumina sequencing were performed to obtain the complete UT1 genome. For nanopore long-read sequencing, high molecular weight genomic DNA was extracted using a Quick-DNA Miniprep kit (Zymo Research) and further purified to remove contaminants using a DNeasy PowerClean Cleanup kit (Qiagen). Libraries were constructed using a Native Barcoding kit and sequenced on a MinION system (Oxford Nanopore Technologies). For Illumina short-read sequencing, genomic DNA was processed for construction of DNA libraries using an Illumina DNA Prep kit and sequenced on the Illumina NovaSeq X plus platform. DNA Sequencing was performed at the University of Florida NexGen DNA Sequencing Core Facility (ICBR; RRID:SCR_019152). Long nanopore reads (12.3 Gb; read length N50 = 9.1 kb) were used for *de novo* assembly using Flye, resulting in one circular contig. The draft genome was then Pilon-polished using 28 million Illumina reads (2x150 bp), which have substantially lower error rates than nanopore reads ^77^. The genome was subsequently annotated, and the phylogenetic tree was built using BV-BRC (https://www.bv-brc.org/). The genome map was plotted using Proksee (https://proksee.ca/). The prophage, ICE/IMEs, bacterial defense and CRISPR-Cas systems were predicted using PHASTEST (https://phastest.ca/), ICEfinder, DefenseFinder, and CRISPRCasFinder (https://crisprcas.i2bc.paris-saclay.fr/CrisprCasFinder/Index), respectively. The CAZymes, CGCs, and their substrates were predicted using dbCAN3 ^38^.

### HPLC analysis

Bacterial supernatants were collected after centrifugation, mixed with H_2_SO_4_ to a final concentration of 4 mM, and then passed through a 0.22 μm filter (EMD Millipore). The concentrations of acetate and butyrate in the supernatants were measured using the Agilent 1220 infinity II LC system composed of an automated sampler, a gradient pump, and a variable wavelength detector. Separations were performed using 4 mM H2SO_4_ as the mobile phase in a fermentation monitoring column (7.8 x 150 mm, Bio-Rad) at 60°C with a flow rate of 0.6 ml/min. The detector wavelength was set at 210 nm. Quantification was based on peak area and the standard curve of SCFAs. Data acquisition and analysis were done using the Agilent ChemStation.

### Mice

C57BL/6 mice were obtained from Jackson Laboratory and maintained under specific pathogen– free conditions. Age-matched (7-8 weeks old) and sex-matched littermates were used in this study. Prior to gavage, female mice were mixed and randomly separated into two groups to minimize potential cage effects. All procedures were conducted in compliance with protocols approved by the Department of Laboratory Animal Resources of the University of Texas Health San Antonio under protocol number 20230051AR.

For preparation of bacterial stocks, single colonies were grown in 500 ml of RCM for ∼24 hours. Cells were harvested by centrifugation, resuspended in anaerobic PBS containing 20% glycerol, aliquoted, and stored at -80°C. For gavaging, bacterial stocks were diluted in anaerobic PBS and transferred to a 1 ml syringe (BD Bioscience) pre-mounted with an animal feeding needle (Fine Science Tools). It was tested that the bacteria retained >95% cell viability after exposing directly the syringe to air for 30 min. A second syringe with anaerobic PBS was prepared in parallel. Both syringes were kept in an airtight AnaeroPouch Bag (Mitsubishi) until animal feeding. Mice were gavaged every 2 days with UT1 (10^9^ CFU in 100 μl PBS), or 100 μl PBS, and fecal samples were collected on day 14. For bacterial colonization, mice were gavaged with a single dose of 10^9^ CFU, and feces were collected on day 0 (baseline), and days 1, 2 and 3 after gavage to isolate genomic DNA. The copy number of *F. prausnitzii* 16S rRNA in the feces was quantified by qPCR using PowerUp SYBR Green Mix (Thermo Fisher), based on the absolute quantification method using a 16S rRNA standard curve ^78^.

For *in vitro* culturing of fecal microbiota, fresh feces were collected from mice gavaged with UT1 (n=9) or PBS (n=5) for 2 weeks (day 14). Feces from each group were pooled and immediately placed in a gut microbiota culture medium (GMM)^79^ (40 mg feces/ml) containing (per liter) 10 g peptone, 10 g beef extract, 3 g yeast extract, 5 g dextrose, 5 g NaCl, 3 g sodium acetate, 0.5 cysteine hydrochloride, 0.4 g NaHCO_3_, 8 mg CaCl_2_, 0.4 mg FeSO_4_, 2 mg MgSO_4_·7H_2_O, 13.6 g KH_2_PO_4_, 0.05 mg vitamin K, 0.1 mg vitamin B12, 0.02 mg biotin, 0.2 mg folate, 0.5 mg riboflavin, 5 mg hemin, and 0.5 mg resazurin (pH 6.7-6.8). The fecal materials were suspended by vortexing and the suspension was allowed to stand at room temperature for 5 min to permit large insoluble particles to settle to the bottom of the tube. The top bacterial suspension was then inoculated into the GMM supplemented with or without soluble starch (1 g per liter) and grown in the anaerobic chamber for measuring OD_600_ values and acetate levels using HPLC.

### 16S rRNA sequencing

Fecal genomic DNA was isolated from the feces of UT1- and PBS-gavaged mice (n = 4 female and 4 males per group) and 16S rRNA libraries were constructed essentially as described previously ^80, 81^. Samples were sequenced to a depth of >92,460 paired-end reads (2 x 300 bp) per sample, which were subsequently processed using QIIM2 (v2022.8). Differences in relative abundance at different taxonomic levels were determined using Student’s *t* test. Linear discriminant analysis effect size (LEfSe) was performed to identify bacterial taxa that significantly contributed to compositional differences at the species level.

### Metagenomic sequencing

Genomic DNA was isolated from the fresh feces collected from UT1- or PBS-gavaged mice (female, n = 4/group) using a ZymoBIOMICS DNA Miniprep Kit (Zymo Research). Note: we used female mice as no sex-dependent microbiome effects were seen based on 16S rRNA analysis. About 500 ng of genomic DNA was used to construct metagenomic DNA libraries using an Illumina DNA Pre kit, which were barcoded using the Illumina DNA/RNA UD indexes. A minimum of 50 million paired-end reads (2 x 150 bp) per sample was obtained. Low-quality sequences of the reads were removed using Trim Galore, and reads mapping to the mouse genome (GRCm38) were then removed using Bowtie2 and Samtools. High-quality microbial reads were mapped to ∼ 1 million bacterial genomes using Metaphlan4 ^82^ to predict the relative abundances of the taxa.

CAZyme mapping and substrate prediction were conducted as described ^83^. Briefly, the microbial reads were assembled into contigs using MEGAHIT v1.2.9, which were passed to Prokka v1.14.6 for gene annotation. Subsequently, the contigs were annotated for CAZymes and CGCs using the dbCAN3 database ^38^ with the protein sequence and gene annotation files as inputs. dbCAN-sub and dbCAN-PUL search approaches were used to predict glycan substrates for CAZymes and CGCs, respectively. The clean reads were then mapped to the nucleotide coding sequences of proteins to calculate the abundance of the CAZymes and CGCs.

### Metabolomic analysis

Two fresh fecal pellets were collected from the same female mice used for metagenomic sequencing as aforementioned and immediately frozen at -80°C for metabolite extraction. Metabolites were analyzed at the Southeast Center for Integrated Metabolomics using a Thermo Q-Exactive Oribtrap mass spectrometer and run in both positive and negative ionization, as described previously ^84, 85^. Metabolite features were normalized to total ion chromatograms, and intensities were tested for group significance using unpaired Student’s *t*-test. Metabolites were identified by comparison to the metabolomic library of purified standards. Metabolic pathway analysis was performed using *Mummichog* v2.6.1. Correlations among the significant bacterial species and metabolites were assessed in the correlation matrix based on Spearman’s rank correlation tests. Matrix was calculated using the rcorr() function of the R package Hmisc and P values were calculated with the rcorr.adjust() function with correction for multiple testing (method “fdr”).

## Statistical analysis

Statistics were performed using unpaired Student *t*-test implemented in GraphPad Prism v10.2.3. p < 0.05 were considered as significant: *, p < 0.05; **, p < 0.01; ***, p < 0.001; Significance in microbe-metabolite correlations was FDR-corrected: *, FDR < 0.05; **, FDR < 0.01.

## Supplemental information

Supplemental information includes supplemental Figures 1 and 2.

## Acknowledgments

We thank Timothy Garrett and Diansy Zincke at the University of Florida for the assistance of LC-MS analysis and DNA sequencing, respectively. This research was supported by start-up support from UT Health San Antonio. R.S.H. is an Investigator of the Howard Hughes Medical Institute and the Ewing Halsell President’s Council Distinguished Chair.

## Author contributions

Y.G. conceived and designed the study. P.G, N.Z. and Y.G. performed experiments and analyzed the data. Y.Z. and R.S.H. contributed to data discussion. Y.G. wrote and revised the manuscript with input from the other authors.

## Data and code availability

Fecal metagenomics raw reads have been made publicly available under the NCBI BioProject accession number PRJNA1130047.

