## Supplementary Figures for "*Faecalibacterium prausnitzii* regulates carbohydrate metabolic functions of the gut microbiome in C57BL/6 mice"

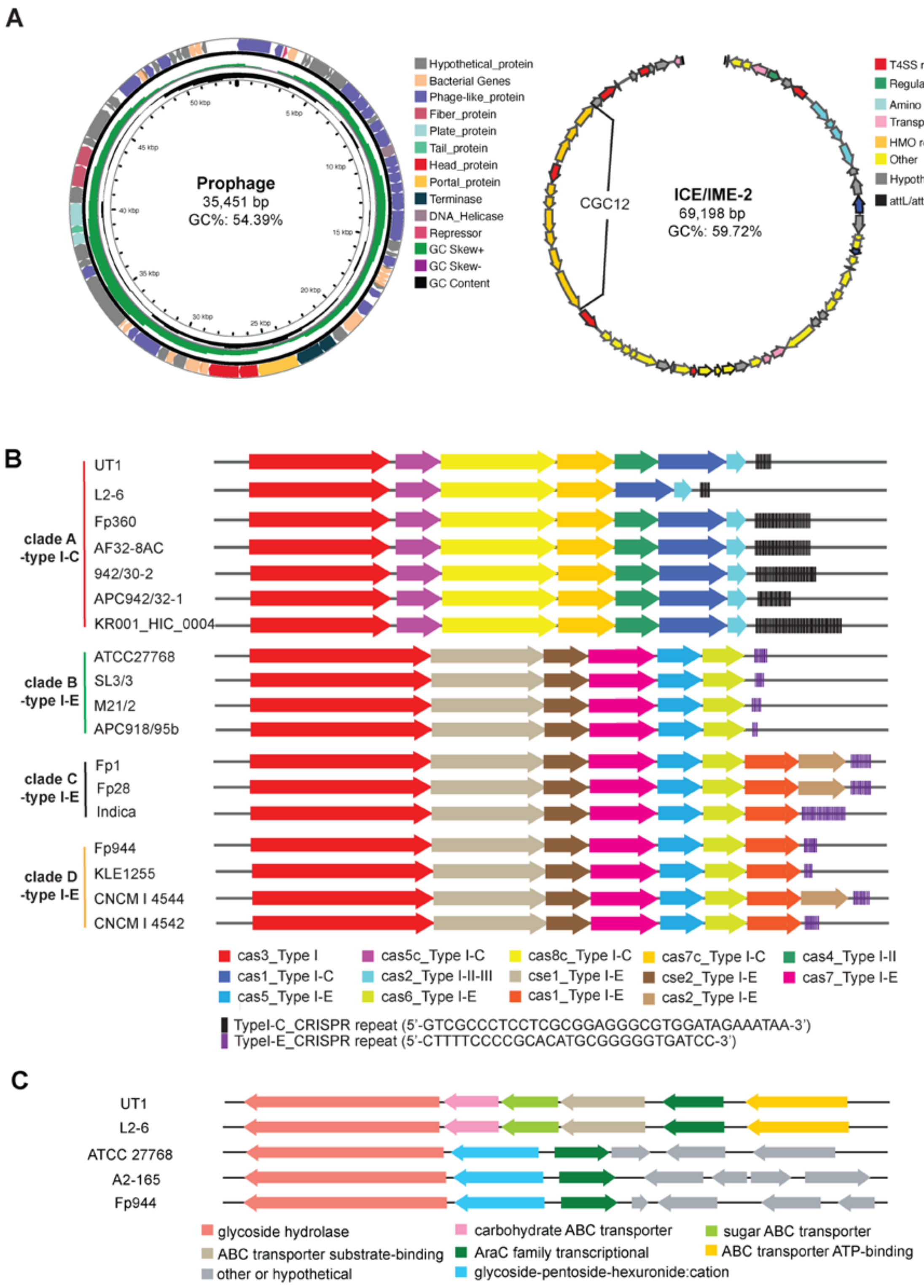


**Supplemental Figure 1. Genetic diversity of *F. prausnitzii* strains. (A)** Genetic maps of the prophage and ICE/IME-2 in UT1 genome**. (B)** CRISPR-Cas systems predicted in strains of *F.*

*prausnitzii* clades A, B, C, and D. **(D)** CAZyme gene clusters (CGCs) predicted to utilize galactomannan among strains of different clades.

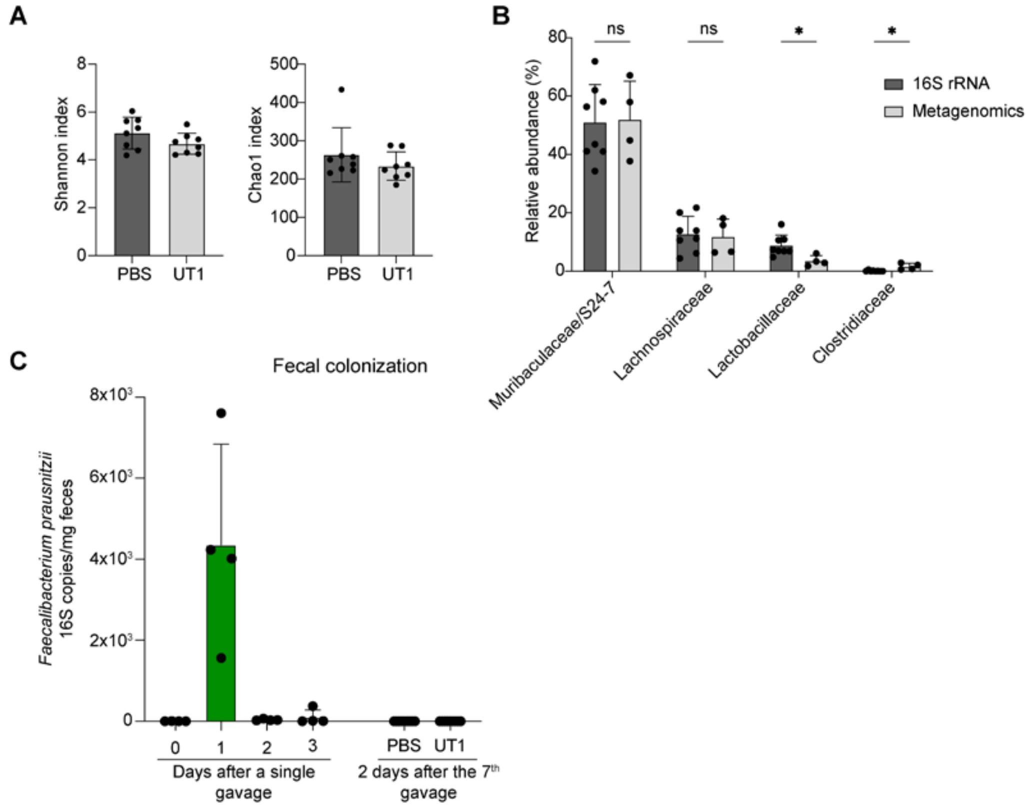


**Supplemental Figure 2. Microbiome diversity of UT1-treated mice. (A)** Shannon and Chao1 indexes, as measurements of alpha-diversity, obtained from 16S rRNA analysis of UT1- and PBS-gavaged mice. **(B)** Comparison of relative abundances of major bacterial families derived from 16S rRNA and metagenomics analyses. **(C)** C57BL/6 mice (n = 4) were gavaged with *F. prausnitzii* strain UT1 ($10^9$ CFU/mouse) one time, and fecal samples were collected every day to detect *F. prausnitzii* 16S rRNA copies by qPCR. The 16S copies were also measured in the fecal samples after 7 gavages on day 14 (n=5/group). *p < 0.05, unpaired student *t* test.
